# Lifespan Extension in Female Mice By Early, Transient Exposure to Adult Female Olfactory Cues

**DOI:** 10.1101/2022.10.07.511218

**Authors:** Michael Garratt, Ilkim Erturk, Roxann Alonzo, Frank Zufall, Trese Leinders-Zufall, Scott D. Pletcher, Richard A. Miller

**Affiliations:** Department of Anatomy, School of Biomedical Sciences, University of Otago, Dunedin, New Zealand; Department of Pathology and Geriatrics Center, University of Michigan, Ann Arbor, MI, USA; Center for Integrative Physiology and Molecular Medicine, Saarland University, 66421 Homburg, Germany; Department of Molecular and Integrative Physiology, University of Michigan, Ann Arbor, MI USA

## Abstract

Several previous lines of research have suggested, indirectly, that mouse lifespan is particularly susceptible to endocrine or nutritional signals in the first few weeks of life, as tested by manipulations of litter size, growth hormone levels, or mutations with effects specifically on early-life growth rate. The pace of early development in mice can also be influenced by exposure of nursing and weanling mice to olfactory cues. In particular, odors of same-sex adult mice can in some circumstances delay maturation. We hypothesized that olfactory information might also have a sex-specific effect on lifespan, and we show here that lifespan of female mice can be increased significantly by odors from adult females administered transiently, i.e. from 3 days until 60 days of age. Female lifespan was not modified by male odors, nor was male lifespan susceptible to odors from adults of either sex. Conditional deletion of the G protein Gαo in the olfactory system, which leads to impaired accessory olfactory system function and blunted reproductive priming responses to male odors in females, did not modify the effect of female odors on female lifespan. Our data provide support for the idea that very young mice are susceptible to influences that can have long-lasting effects on disease resistance, and provide the first example of lifespan extension by olfactory cues in mice.

## Introduction

Interventions in the first few weeks of postnatal life seem to have particular power to produce life-long changes in mouse physiology, with effects on late-life illnesses and on survival. Bartke’s laboratory, for example, has shown that the exceptionally long lifespan of the Ames dwarf mice, in which longevity appears to reflect lower levels of growth hormone (GH) produced by the anterior pituitary, can be reduced back to that of non-mutant controls by transient exposure to daily GH injections started at 2 weeks of age and discontinued 6 weeks later [1]. Furthermore, these early-life GH injections block the development of multi-modal stress resistance shown by skin-derived fibroblasts of Ames dwarf mice [1], and also prevent the lower levels of hypothalamic inflammatory characteristic of unmanipulated Ames mice [2]. Many other characteristics of Ames dwarf mice, plausibly connected to their longevity, insulin sensitivity [3], and cognitive health [4, 5], are also blocked by early-life GH treatments when tested at 18 months of age, including changes in fat, muscle, plasma irisin and GPLD1, liver, and brain [6]. Conversely, lifespan can be increased in genetically normal mice by pre-weaning manipulation of litter size – mice weaned from litters in which litter size has been increased to 12 pups/nursing mother live longer than those in control litters of size 8 [7]. Genetic selection experiments have also provided relevant data, by showing extended lifespan in non-inbred stocks of mice that were selected, for 17 or more generations, for exceptionally slow growth in the first 10 days of life, or between 26 – 56 days of age [8].

Age at sexual maturity is another early-life developmental factor that has been associated with lifespan. Across strains of mice, age at sexual maturity correlates positively with median lifespan [9], and genetic manipulations that delay sexual maturity have been associated with longer lifespan in females [10]. Age at sexual maturity in rodents is also influenced by environmental cues, in particular olfactory cues from con-specifics. Female mice that are exposed to male odors (typically soiled bedding or male urine) during development show hastened sexual maturity [11, 12]. In contrast, exposing females to the odors of group-housed adult virgin females can delay sexual maturity [12, 13]. In males, exposure to females or their odors has been shown to be associated with increased masses of reproductive organs (testes, seminal vesicles) early in life, also indicating earlier sexual maturity in males [14]. These odor priming effects are assumed to be adaptive, allowing animals to modulate their timing of early life reproduction to suit predicted future environments. In one previous study, hastened female sexual maturity was associated with an increase in mortality over the first 180 days of life, suggesting a cost to this developmental change [15]. However, in this prior study females were exposed directly to males on the day of their first estrous, which was earlier in females exposed to male odors producing a confounding effect that could contribute to mortality differences.

The effects of odors on sexual maturity are initiated, at least partly, by detection of odorants through the vomeronasal organ, one of the two olfactory sub-systems in mice. Female mice that have had their vomeronasal organ surgically removed do not show hastened sexual maturity in response to male odors [16]. Vomeronasal sensory neurons (VSNs) express hundreds of different sensory receptors, but these receptors largely use one of two different types of G-protein for signal transduction [17]. The G protein Gαo is used for signal transduction in VSNs of the basal layer of the vomeronasal organ. Cre-mediated deletion of Gαo in the olfactory system has been shown to inhibit females from showing hastened sexual maturity in response to male odors [18], indicating that this subset of VSNs is important in driving changes in sexual maturity, at least in response to opposite sex cues in females.

With these precedents in mind, we tested the idea that olfactory signals, presented to mice from 3 days of age but discontinued at 60 days, would have long-lasting effects that influence age at death. We initiated odor exposure early in life, prior to weaning, because a previous study had shown that very early exposure influences female body weight during development [19]. In particular, we speculated that odor from adult female mice would lead to lifespan extension in female (but not in male) mice, and vice versa. We also hypothesized that exposure to opposite-sex odors would reduce survival given the hastening of sexual maturity. Here we show that transient exposure to odor from adult females starting at 2 days of age can extend lifespan of female mice.

## Results

### Survival

Our principal hypothesis was that early-life exposure to sex-specific olfactory cues indicative of social environment would influence lifespan of mice. A secondary hypothesis was that the response to those cues would depend on function of Gao-expressing neurons in the vomeronasal organ, since these have been shown to mediate early life priming responses to opposite-sex urine, at least in female mice [18]. Mice were exposed either to same-sex or to opposite-sex odors starting from three days of age, with daily exposure through day 60 of life. They were housed in same-sex cages from the age of weaning, i.e. from 19 or 20 days. **Figure 1** shows survival curves for female and for male mice subjected either to odors from adult males (“MU”) or from adult females (“FU”), or exposed to water as a control (ZU, for zero urine). Data from male and female mice were evaluated separately. **Table 1** shows median age, and age at 90^th^ percentile, for each combination of Sex and Treatment, pooling across genotypes. In female mice, FU led to an 8% increase in median lifespan, compared to the ZU control group, and a 9% increase in the age at 90^th^ percentile. The log-rank test, comparing all three treatment groups, showed that treatment led to significant differences among the groups (p = 0.04), and a follow-up test showed that for females the FU mice were longer-lived than the ZU controls (p = 0.01 by log-rank test). Male odors did not produce a lifespan change in female mice (p = 0.6). Treatment did not modify lifespan in the male mice; log-rank p = 0.8 for all treatment groups taken together).

**Figure 1:**
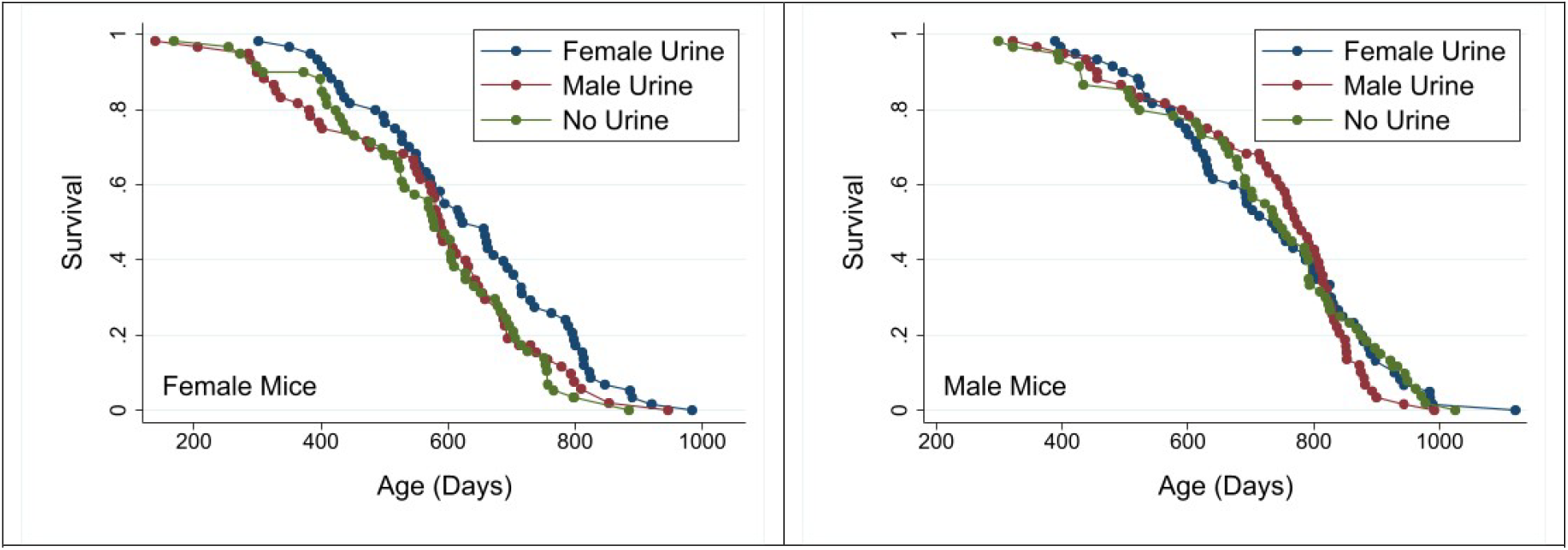
survival curves for female (A) and male (B) mice exposed to male, female, or no odors daily from 2 days to 60 days of age. Each dot represents the age at death of an individual mouse. Both genotypes of mice are pooled for analysis. See the Supplementary Data for age at death of each mouse and its genotype, sex and treatment group.

**Table 1:**
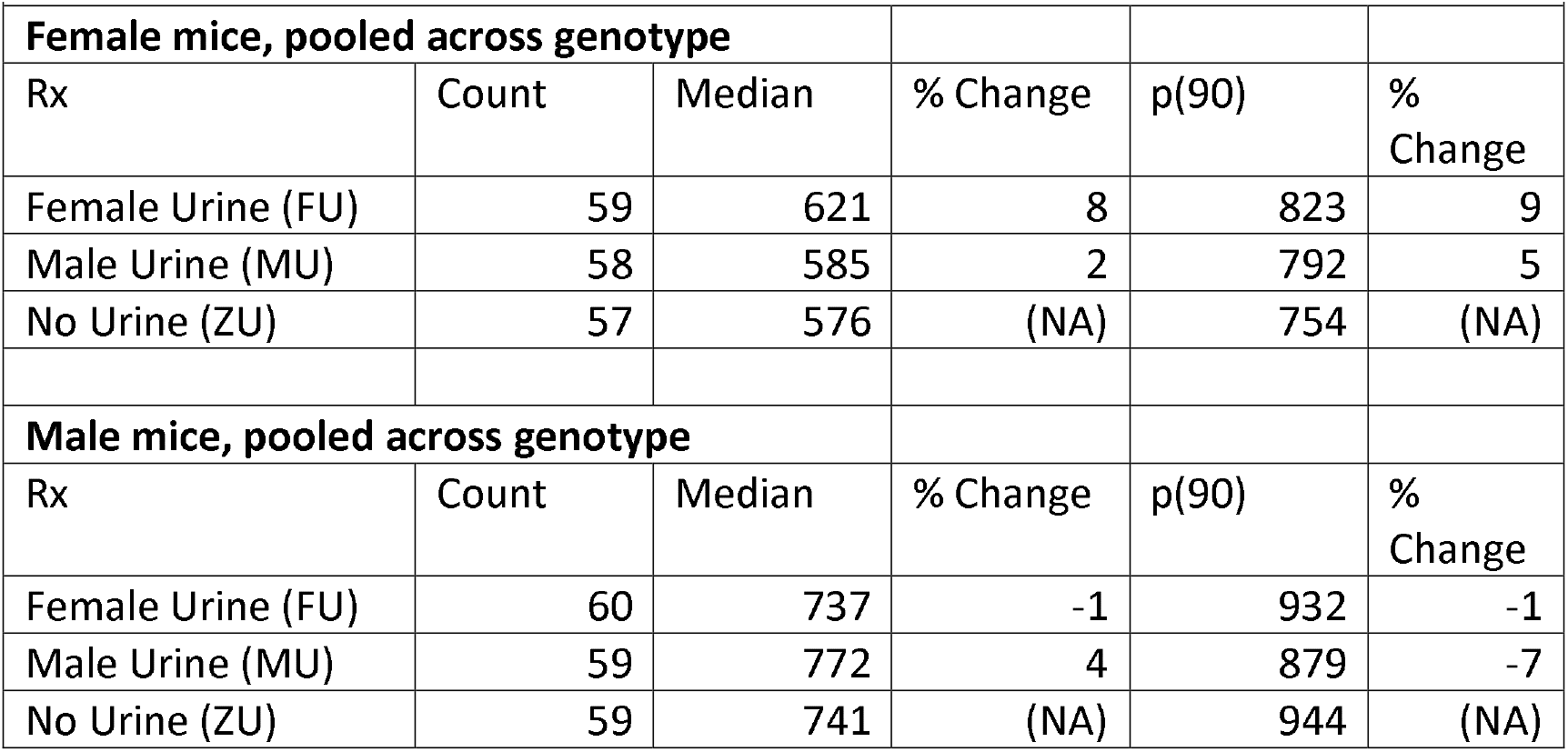
Median and 90^th^ Percentile Survival Statistics for Mice Treated With Same-Sex or Opposite-Sex Odors from 2 to 60 Days of Age.

|  |  |  |  |  |  |
| --- | --- | --- | --- | --- | --- |
| Female mice, pooled across genotype |  |  |  |  |  |
| Rx | Count | Median | % Change | p(90) | % Change |
| Female Urine (FU) | 59 | 621 | 8 | 823 | 9 |
| Male Urine (MU) | 58 | 585 | 2 | 792 | 5 |
| No Urine (ZU) | 57 | 576 | (NA) | 754 | (NA) |
| Male mice, pooled across genotype |  |  |  |  |  |
| Rx | Count | Median | % Change | p(90) | % Change |
| Female Urine (FU) | 60 | 737 | -1 | 932 | -1 |
| Male Urine (MU) | 59 | 772 | 4 | 879 | -7 |
| No Urine (ZU) | 59 | 741 | (NA) | 944 | (NA) |

To see if the effect of odor was dependent on Gao expression in the vomeronasal organ, survival data were analyzed by Cox regression, with two factors: Treatment (MU, FU or ZU), and Genotype (WT for wild-type (Gnao1^fx/fx^, OMP-cre -/-), or OMP-Mutant (Gnao1^fx/fx^, OMP-cre +/-)), with a [Treatment x Genotype] interaction term. Table 2 shows the significance levels for each of the three terms, calculated separately for female and for male mice. The interaction term did not achieve significance in either sex, implying that the odor effect did not depend on genotype. In addition, there was no independent effect of genotype on survival. Only the odor treatment had a significant effect on survival, and only in female mice (p = 0.04), consistent with the results of the log-rank tests. The Cox regression calculation also confirmed the inference that FU females lived longer than ZU females (p = 0.015 without adjustment for multiple comparisons, and p = 0.045 with Sidak adjustment), and that no other pairwise comparison between treatment groups had a significant effect in either sex.

**Table 2:**
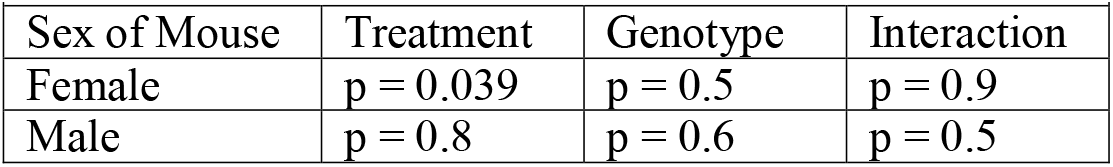
Significance Tests for Predictors in Cox Regression, Calculated Separately for Each Sex.

| Sex of Mouse | Treatment | Genotype | Interaction |
| --- | --- | --- | --- |
| Female | $p = 0.039$ | $p = 0.5$ | $p = 0.9$ |
| Male | $p = 0.8$ | $p = 0.6$ | $p = 0.5$ |

To evaluate the effects of early-life odor exposure on survival to unusually high ages, we used the method of Wang and Allison [20], which calculates a Fisher Exact test statistic on the proportion of mice in each treatment that remain alive at the 90^th^ percentile of their joint survival distribution, the method also employed by the mouse Intervention Testing Program consortium [21]. In females, the comparison of FU to control ZU mice produced p = 0.004, showing that FU-treated females were more likely to reach old age than ZU-treated females. None of the other paired comparisons reached significance in female mice, and none did so in male mice.

### Functional testing

Each mouse surviving to 22 months of age was evaluated using a series of tests for age-sensitive physiological function. The data were analyzed separately for each sex, using a two-factor ANOVA (Treatment, Genotype, Interaction). There were no treatment effects, or significant interaction terms, for grip strength (front paws or four paws, mean or maximum of three trials) or for rotarod performance (mean or maximum of three trials) in either sex. There was, however, an effect of early-life odor exposure on core body temperature, seen in females only (p = 0.01), and a Sidak post-hoc test showed that body temperature was higher in FU females than in ZU controls (p = 0.007). **Figure 2a** shows the dotplots for female mice in the three treatment groups. Since core body temperature declines in age in mice, this result is consistent with the lifespan benefit noted in the FU vs ZU comparison. Male mice did not show any effect on body temperature (not shown). Weight was measured at 1, 2, 3, 6, 12, 18, and 24 months of age, and there were no differences among treatment groups, except at 6 months of age (**Figure 2b**), at which ZU females were marginally higher in weight than mice in the MU or FU groups (p = 0.03; if the data are tested without the MU outlier weighing 16 grams, p = 0.04). Plasma glucose was measured in non-fasting animals at 12 months of age. There were no effects of treatment in female mice, but glucose levels were significantly lower in male ZU mice than in the FU or MU mice (**Figure 2c**; p = 0.004 for ANOVA, and p = 0.01 for each pairwise comparison to ZU male mice.)

**Figure 2:**
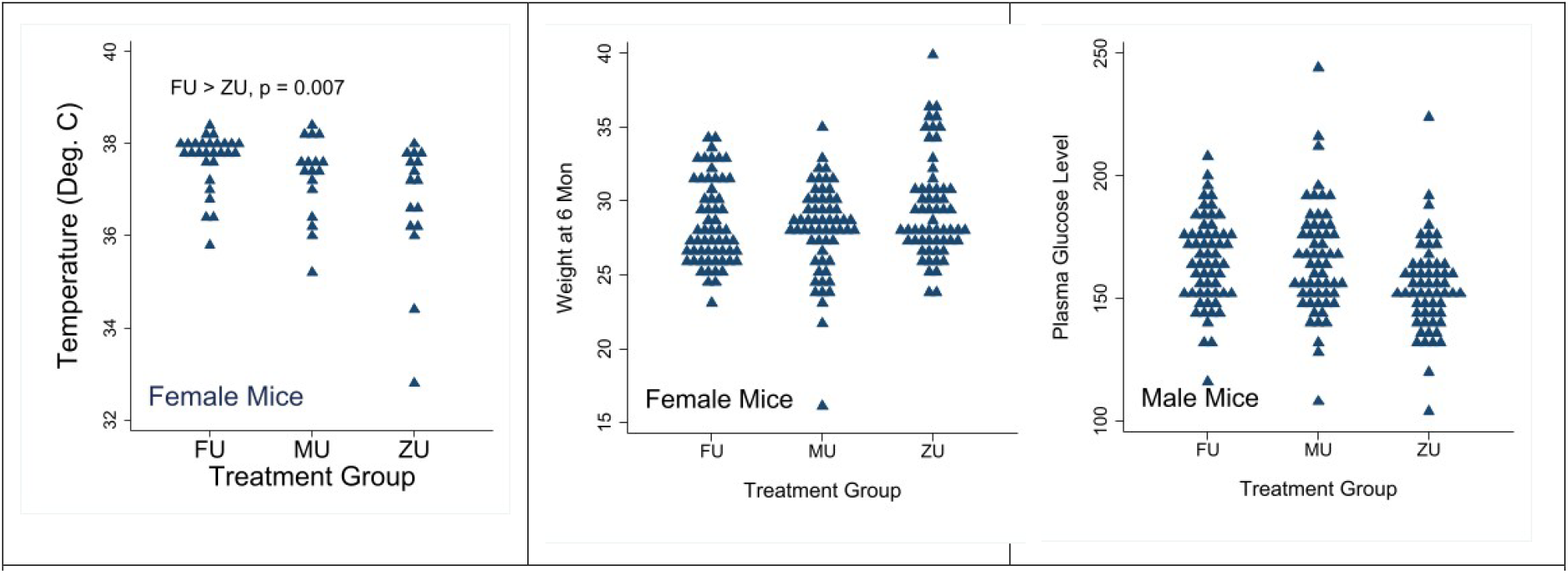
(a) Core body temperature (°C) in female mice at age 22 months of age in each odor-treatment group. (b) Weight (g) at 6 months of age in female mice. (c) Plasma glucose (mg/dl) in male mice at 12 months. Each symbol represents the value for an individual mouse. Data is pooled for genotypes, with full data matched to each genotype and treatment group available in the supplementary Data.

## Discussion

Our results add to the growing evidence that late life health, including risks of lethal diseases, can be influenced by non-genetic factors, such as litter size [7, 22] and GH levels [1, 2], to which a newborn or pre-weanling mouse is exposed transiently. The “window of opportunity,” or age at which a mouse is susceptible to such influences, is not known, and may differ depending on the specific form of environmental variation, but it is noteworthy that GH treatments can prevent lifespan extension of Ames dwarf mice when these are initiated at two weeks of age, but fail in Snell dwarf [23] and Ames dwarf animals (Bartke, unpublished results) when started at four weeks of age. The basis for this transient susceptibility to factors that set the tempo of late-life decline is unknown, but seems likely to involve epigenetic modulation of events that are sensitive to nutritional and neuro-endocrine stimuli. Fibroblast cultures from adult mice of the Ames and Snell dwarf mutant strains are resistant, in culture, to multiple forms of lethal injury [24–26], unless the Ames mice had been subjected to early-life GH treatment [1]. In this context it is noteworthy that fibroblasts derived from one-week old Snell dwarf and control mice do not differ in stress resistance [25]; at this age GH levels are still largely derived from the intra-uterine blood circulation, and are not yet controlled by the pituitary of the newborn mouse itself. GH-exposure experience in the 2 – 8 week old mouse [2], as well as milk availability mediated by the size of the nursing litter [22] also modulate the hypothalamic inflammatory state in mice 18 months of age or older. Ames mice treated transiently with GH also resemble wild-type, i.e. non-mutant, controls in adipose tissue UCP1 levels, relative ratios of pro-inflammatory and anti-inflammatory macrophages in adipose tissue, production of FNDC5 and its myokine cleavage product irisin by skeletal muscle, hepatic and plasma levels of GPLD1, and brain levels of protective factors BDNF and DCX [6], showing that these early-life environmental factors can have long-lasting, presumably permanent, effects of many physiological properties of high relevance to health maintenance and disease resistance.

The data reported here document another vector for early-life modulation of late-life fitness and mortality risks, by manipulation of con-specific social odors. We chose to study this topic because of data showing effects of early exposure to same-sex and opposite-sex odors on the pace of mouse development and sexual maturation. We hypothesized that exposure to same-sex odor, which delays development in some circumstances [15, 27], might also delay the rate of age-related decline, and we found that indeed female mice exposed early to odors from adult females had longer lifespan than females exposed to water. Male odor did not have this effect on female longevity. The FU-treated female mice also resisted the age-related decline in core body temperature, although there was no inter-group difference in grip strength or rotarod performance, other indices of late-life physiological state. Male mice, in contrast, did not exhibit lifespan or performance effects in response to either same-sex or opposite-sex odors. We do not know the mechanism for the lifespan effect in the FU-treated female mice, or why we saw no such effect in male animals; these are questions we plan to explore in future work.

The effects of odor exposure on lifespan were not modified by deletion of Gαo in the olfactory system. This G protein is important in mediating responses to peptide and protein-based ligands in basal VSNs [28], and has previously been shown to inhibit priming responses to male odours in females, including changes in sexual maturity [18]. We therefore designed this experiment using this model because this deletion is known to inhibit priming effects, but does not affect normal body development or suckling behaviour as has been observed with manipulations that impair main olfactory system function [29]. Previously published research with this model has largely focused on responses to male odours, but in this study we observed changes in lifespan in response to female odours. Exposure of female mice to female odours has previously been shown to activate a different subset of VSNs compared to male odorants, including apical VSNs that use a different G protein in signal transduction [30]. Thus, effects of female odors on female survival may be mediated by a subset of VSNs different from those that were inhibited in this study. It is also possible that effects could be mediated by detection of odorants by the main olfactory system, or a different sensory modality, since females were exposed to a mixture of odors and other cues that would be found in soiled bedding. Changes in sexual maturity with exposure to conspecific odors can be initiated even by brief exposure to the diluted urine of a conspecific after weaning, although longer term effects on female body weight have been observed when odor exposure begins prior to weaning [31], which is why we chose an extended treatment protocol to maximise the chance of detecting effects on survival. Future studies with a shorter treatment time course after weaning, and with only urine or specific urinary fractions, would help to reveal the signals and developmental periods that important causing subsequent changes in survival.

Studies of the effects of sensory perception on aging and lifespan in invertebrates dates back at least to the work of Apfeld and Kenyon in Caenorhabditis elegans [32], and in the years since, a variety of sensory modalities, including smell, taste, sight, and pain, have become established as important modulators of aging across invertebrate taxa [33–35]. Exposure of flies and worms to food-based odorants, for example, limit the benefits of dietary restriction and influence measures of healthy aging, including sleep and daily activity patterns [36, 37]. Some of these studies have focused on how perception of conspecific pheromones, detected through olfaction and gustation, can modulate Drosophila and C. elegans lifespan [38–40]. Although there is no guarantee that mechanisms linking pheromones to aging in worms and flies will prove to be analogous to pathways in mice, they do provide clues that deserve to be explored. Worm perception impacts insulin-specific pathways, and conserved motivation and reward neuropeptides, such as NPF/NPY, are required for aging effects in flies [38–40]. It seems reasonable to test whether these same pathways influence mammalian lifespan in mice, either with, or in the absence of, sensory cues of the kind revealed by our experimental findings.

As far as we know, this is the first observation that lifespan can be increased, in a mammal, by olfactory signals. More generally, the work hints that very young mice, and perhaps animals of other species, are able to assess aspects of the social environment and adjust aspects of their development that might improve their Darwinian fitness and, as a side-effect, slow the rate of aging and make them more resistant to late-life disease.

## Methods

### Mice

The proposed lifespan experiment was approved by the University of Michigan Animal Care and Use Program, Protocol PRO00007884. All experiments strictly adhered to this approved protocol. All mice contained a version of the G□o gene (Gnao1) that had been floxed at exons five and six. These were crossed with mice carrying a transgene of Cre recombinase that caused cre expression under the control of olfactory marker protein (*OMP*^cre^). This cross has been shown [28] to inhibit G□o expression specifically within vomeronasal sensory neurons (VSNs). The background stock of this cross is a mix of the C57BL/6J and S129 mouse strains. All mice used in this study were the offspring of parents that were both homozygous for the floxed G□o gene, with one parent in each pair (either the mother or the father) also containing one copy of the transgene for Cre recombinase. This mating system led to the production of Cre positive mice (Gnao1^fx/fx^, OMP-cre +/-, with deletion of G□o in VSNs) and cre negative (Gnao1^fx/fx^, OMP-cre -/-, control) mice in equal proportions. Mice in each individual litter were randomly assigned to a specific treatment arm (male odor, female odor, or no odor, see below) at age 2 days, and were weaned to same-sex cages at 4 mice/cage at 19 or 20 days of age. Litters of size 7 or lower were not used, and litters with more than 8 pups were trimmed to 8 pups as soon as discovered. A tail snip biopsy was taken for genotyping at age 10 – 19 days. Each cage of weanlings contained four mice, two wild-type and two mutant, and each mouse in a cage was in the same odor-exposure treatment group. Cages were labeled with details of mouse sex, genotype and treatment group and so were not blinded, although the technician team that assessed mice on a daily basis were largely unaware of the principal hypotheses of the proposed experiment. Thirty mice were allocated per sex, genotype and odor treatment combination with each mouse assumed to be an independent biological replicate. This sample size was selected because it provides sufficient power to detect a change in median lifespan of approximately 10-15% within one sex and genotype. We originally hypothesized that some effects would be consistent across sexes allowing sexes to be pooled for analysis, further increasing statistical power. All mice received Purina 5LG6 food and water without restriction.

### Odor exposure

Urine was collected from group-housed (3 males or 4 females per cage) young adult (6 month old) male or female UM-HET3 mice [41], and stored in aliquots at −20°C until use. Aliquots were then thawed when needed and discarded within 24 hr of thawing; no aliquot was used on more than a single day. Newborn mice had their noses moistened, using a cotton swab, with female urine (FU), male urine (MU), or autoclaved water (ZU, control) six days each week, starting at Day 3 of life, where Day 1 is the date on which pups were first noted in the breeding cage. Urine was applied in the light period between 8 am and noon each day. The mother was placed in a clean cage just prior to odor-exposure of her pups, and then returned to the nursing cage immediately thereafter.

After weaning, mice were exposed to same-sex, opposite-sex, or control soiled bedding instead of urine. Spent bedding was obtained from cages housing 4 adult female or 3 adult male UM-HET3 mice, aged 4 – 12 months, after the cage had been occupied for seven days. Bedding from each donor cage was thoroughly mixed by hand, and then 10% of bedding from a donor cage was placed in each recipient cage, with an equal proportion of the bedding from the recipient’s cage removed. Spent bedding was added once each day for six days per week until mice were 12 weeks of age. For control cages, no bedding was added but the bedding in the cage was re-arranged to simulate the MU and FU intervention.

### Lifespan data

Lifespan analysis and criteria for inclusion in final survival statistics followed the established protocols of the Interventions Testing Program [21]. Each mouse was inspected daily and the date of natural death recorded. Mice were euthanized to comply with humane-use protocols if they appeared unlikely to survive more than another 7 days using a symptom checklist. Animals were excluded from analysis and censored if they were injured due to fighting, showed malocclusion, or had dermatitis covering greater than 20% of their body. Six females and two males were removed from the study as a consequence of these exclusion factors, leaving the final sample sizes for survival analysis presented in Table 1.

### Assessment of temperature, grip strength, and rotarod performance

Mice were evaluated at 22 months of age. Cages of mice were transferred to the testing room between 8 am and 10 am, and immediately evaluated for core body temperature using a rectal thermometer (Braintree RET3). Mice in the cage were then individually weighed. The mice were then allowed to acclimate in their home cage for one hour prior to further testing, after which grip strength was evaluated. Mice were held by the base of the tail and lowered toward a wire grid connected to a strain gauge (Bioseb Research Instruments). Once both forepaws grasped the grid, they were firmly but gently pulled horizontally until they released their grip. Strain gauge measurements were recorded for each of three trials for each mouse (“forepaw grip”). Three additional trials were then conducted, allowing all four paws to simultaneously grasp the grid (“allpaws grip”). The cage was then returned to its original housing room. On the next day, cages were again moved to the testing room and mice allowed to acclimate for an hour, after which a RotaRod test device (Ugo Basile) was used to measure motor coordination on an accelerating, rotating rod, for three trials. Mice were placed on the RotaRod at an initial speed of 5 RPM. The Rotarod accelerated to a maximum of 40 RPM over 300 seconds. The trial ended when the mouse fell from the RotaRod, and the latency to fall was recorded. Each mouse was tested three times, with a one minute interval between trials to allow the apparatus to be cleaned.

### Assessment of non-fasting blood glucose and weight

Blood samples (approximately 50 microliters) were taken by tail venipuncture from restrained, non-anesthetized 12 month old mice into 300 microliter tubes containing EDTA, and tested using a Bayer Contour Next EZ Glucometer. Body weight was periodically assessed between 9 am and noon.

### Statistical analysis

We tested for differences among treatment groups, for each sex, using the log-rank test, with the two genotypes pooled. When the overall log-rank test, with all three odor groups, failed to confirm the null hypothesis, we then proceeded to consider all pairs of odor groups using the log-rank test. To determine if genotype modulated the response to odors, we used Cox regression with two factors, genotype and odor group, including the interaction term, and did so for each sex separately. The significance of the interaction term was used as the indicator of whether response to odor was or was not genotype-dependent within each sex. As a surrogate for “maximum” lifespan, we used the Wang/Allison test [20], comparing odor groups for the proportion of mice still alive at the 90^th^ percentile of the joint survival distribution. Calculations of median and 90^th^ percentile age were performed on the number of mice shown in Table 1, i.e. excluding mice that had died because of fighting or other accident. All analyses were conducted using STATA version 17.

## Data Availability

All survival data and data used for figure two are provided as supplementary information.

## Acknowledgements

This work was supported by NIA grant AG024824 and by the Glenn Foundation for Medical Research. Funding from the American Federation for Aging Research, The Michigan Society of Fellows, and The Marsden Fund are also acknowledged.

